# Unsupervised Learning of Persistent and Sequential Activity

**DOI:** 10.1101/414813

**Authors:** Ulises Pereira, Nicolas Brunel

**Affiliations:** Department of Statistics, The University of Chicago, Chicago, Illinois, United States of America; Department of Neurobiology, The University of Chicago, Chicago, Illinois, United States of America; Department of Neurobiology, Duke University, Durham, North Carolina, United States of America; Department of Physics, Duke University, Durham, North Carolina, United States of America

**Keywords:** unsupervised learning, persistent activity, sequential activity, synaptic plasticity, Hebbian plasticity, homeostatic plasticity

## Abstract

Two strikingly distinct types of activity have been observed in various brain structures during delay periods of delayed response tasks: Persistent activity (PA), in which a sub-population of neurons maintains an elevated firing rate throughout an entire delay period; and Sequential activity (SA), in which sub-populations of neurons are activated sequentially in time. It has been hypothesized that both types of dynamics can be ‘learned’ by the relevant networks from the statistics of their inputs, thanks to mechanisms of synaptic plasticity. However, the necessary conditions for a synaptic plasticity rule and input statistics to learn these two types of dynamics in a stable fashion are still unclear. In particular, it is unclear whether a single learning rule is able to learn both types of activity patterns, depending on the statistics of the inputs driving the network. Here, we first characterize the complete bifurcation diagram of a firing rate model of multiple excitatory populations with an inhibitory mechanism, as a function of the parameters characterizing its connectivity. We then investigate how an unsupervised temporally asymmetric Hebbian plasticity rule shapes the dynamics of the network. Consistent with previous studies, we find that for stable learning of PA and SA, an additional stabilization mechanism, such as multiplicative homeostatic plasticity, is necessary. Using the bifurcation diagram derived for fixed connectivity, we study analytically the temporal evolution and the steady state of the learned recurrent architecture as a function of parameters characterizing the external inputs. Slow changing stimuli lead to PA, while fast changing stimuli lead to SA. Our network model shows how a network with plastic synapses can stably and flexibly learn PA and SA in an unsupervised manner.

## INTRODUCTION

Selective persistent activity (PA) has been observed in many neurophysiological experiments in primates performing delayed response tasks, in which the identity or spatial location of a stimulus must be maintained in working memory, in multiple cortical areas, including areas in the temporal lobe (Fuster et al., 1982; Miyashita, 1988; Miyashita and Chang, 1988; Sakai and Miyashita, 1991; Nakamura and Kubota, 1995; Naya et al., 1996; Miller et al., 1996a; Erickson and Desimone, 1999), parietal cortex (Koch and Fuster, 1989; Chafee and Goldman-Rakic, 1998) and prefrontal cortex (Fuster et al., 1971; Funahashi et al., 1989, 1990, 1991; Miller et al., 1996b). More recently, selective persistent activity has also been observed in mice (Liu et al., 2014; Guo et al., 2014; Inagaki et al., 2017) as well as flies (Kim et al., 2017). It has been hypothesized that PA represents the mechanism at a network level of the ability to hold an item in working (*active*) memory for several seconds for behavioral demands. Theoretical studies support the hypothesis that persistent activity is caused by recurrent excitatory connections in networks of heavily interconnected populations of neurons (Amit et al., 1994; Durstewitz et al., 2000; Wang, 2001; Brunel, 2005). In these models, PA is represented as a fixed point attractor of the dynamics of a network that has multiple stable fixed points. The connectivity matrix in such models has a strong degree of symmetry, with strong recurrent connections between sub-groups of neurons which are activated by the same stimulus. This connectivity matrix can be learned by modifying recurrent connections in a network according to an unsupervised Hebbian learning rule (Mongillo et al., 2005; Litwin-Kumar and Doiron, 2014; Zenke et al., 2015).

Sequential activity (SA) has been also observed across multiples species in a number of behaviors such as spatial navigation (Foster and Wilson, 2006; Harvey et al., 2012; Grosmark and Buzsáki, 2016) and bird song generation (Hahnloser et al., 2002; Amador et al., 2013; Okubo et al., 2015). Furthermore, a large body of experimental evidence shows that SA can be learned throughout experience (Okubo et al., 2015; Grosmark and Buzsáki, 2016). Several theoretical network models have been able to produce SA (Abeles, 1991; Amari, 1972; Kleinfeld and Sompolinsky, 1988; Diesmann et al., 1999; Izhikevich, 2006; Liu and Buonomano, 2009; Fiete et al., 2010; Waddington et al., 2012; Cannon et al., 2015). In these models, the connectivity contains a feed-forward structure - neurons active at a given time in the sequence project in a feed-forward manner to the group of neurons which are active next. From a theoretical stand point, the mechanism to generate SA is fundamentally different from the one that generates PA. While SA usually corresponds to a path in the state space of the network, PA is identified as a fixed point attractor. Thus, SA has an inherent transient nature while PA is at least linearly stable in a dynamical system sense.

The question of how sequential activity can be learned in networks with plastic synapses has received increased interest in recent years. The models investigated can be roughly divided in two categories: models with supervised and unsupervised plasticity rules. In models with supervised plasticity rules, the synapses are updated according the activity of the network and an *error signal* that carries information about the difference between the current network dynamics and the one that it is expected to learn by the network (Sussillo and Abbott, 2009; Memmesheimer et al., 2014; Laje and Buonomano, 2013; Rajan et al., 2016). In models with unsupervised plasticity rules, sequential dynamics is shaped by external stimulation without an error signal (Jun and Jin, 2007; Liu and Buonomano, 2009; Fiete et al., 2010; Waddington et al., 2012; Okubo et al., 2015; Veliz-Cuba et al., 2015). In those models SA is generated spontaneously, and the temporal statistics of the stimulation shapes the specific timing of the sequences.

Both experimental and theoretical work therefore suggest that neural networks in the brain are capable to learn PA and SA. One unresolved issue is whether the learning rules used by brain networks to learn PA are fundamentally different than the ones used to learn SA, or whether the same learning rule can produce both, depending on the statistics of the inputs to the network. Learning rules employed in theoretical studies to learn PA typically do not contain any temporal asymmetry, while rules used to learn SA need to contain such a temporal asymmetry.

Here, we hypothesize that a single learning rule is able to learn both, depending on the statistics of the inputs. We investigate what are the conditions for the plasticity mechanisms and external stimulation to learn PA or SA using unsupervised plasticity rules. We consider a model composed of multiple populations of excitatory neurons, each activated by a distinct stimulus. We consider a sequential stimulation protocol in which each population of neurons is stimulated one at a time, one after the other. This protocol is characterized by two parameters, the duration of stimulus presentations and the time interval between stimulations. This simple setting allows us to explore between the extremes of isolated stimulations with short or large duration and sequential stimulations close or far apart temporally. We use a rate model to describe the activity of populations of neurons (Wilson and Cowan, 1972). The connectivity in this model represents the average of the synaptic connections between populations of neurons, allowing to investigate at a mesoscopic level the learning mechanisms of PA and SA. This model has the advantage of analytical tractability.

This paper is organized as follows: We first characterize the types of possible dynamics observed in network with both feed-forward and recurrent connections, in the space of possible (fixed) connectivities. We then show that a network with plastic connections described by a unsupervised temporally asymmetric Hebbian plasticity rule stimulated sequentially does not stably learn PA and SA. We then explore two types of stabilization mechanisms: 1) synaptic normalization; 2) a multiplicative learning rule. We show that when a synaptic normalization mechanism is included, PA and SA cannot be learned stably during sequential stimulation. However, the addition of a modified multiplicative learning rule leads to successful learning of PA or SA, depending on the temporal parameters of external inputs, and the learning can be characterized analytically as a dynamical system in the space of fixed connectivities parametrized by the stimulus parameters.

## 1 METHODS

### 1.1 Networks with fixed connectivity

We first consider three different *n* population rate models that share in common two connectivity motifs that have been classically considered a distinctive feature of PA and SA respectively: recurrent and feed-forward connections. The three network models considered are: 1) *n* excitatory neurons; 2) *n* excitatory neurons with shared inhibition; 3) *n* excitatory neurons with adaptation. The strength of the recurrent and feed-forward connections are *w* and *s* respectively. We used the current based version of the widely used firing rate model, which is equivalent to its rate based version (Miller and Fumarola, 2012) with three different nonlinear transfer functions.

#### 1.1.1 Network of excitatory neurons

The network consists in *n* excitatory populations connected by feed-forward and recurrent connections with strength *w* and *s* respectively as it is shown in Fig 1A.I. The dynamics is given by:

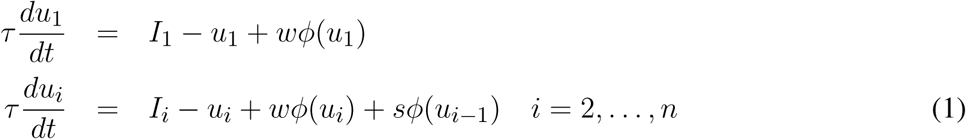

where *I*_*i*_ represents the external input to neuron *i, τ* is the characteristic time scale for excitatory populations and *ϕ*(*u*) is the current to average firing rate transfer function (or f-I curve). The resulting average firing rates are denoted by *r*_*i*_ *≡ ϕ*(*u*_*i*_).

**Figure 1.**
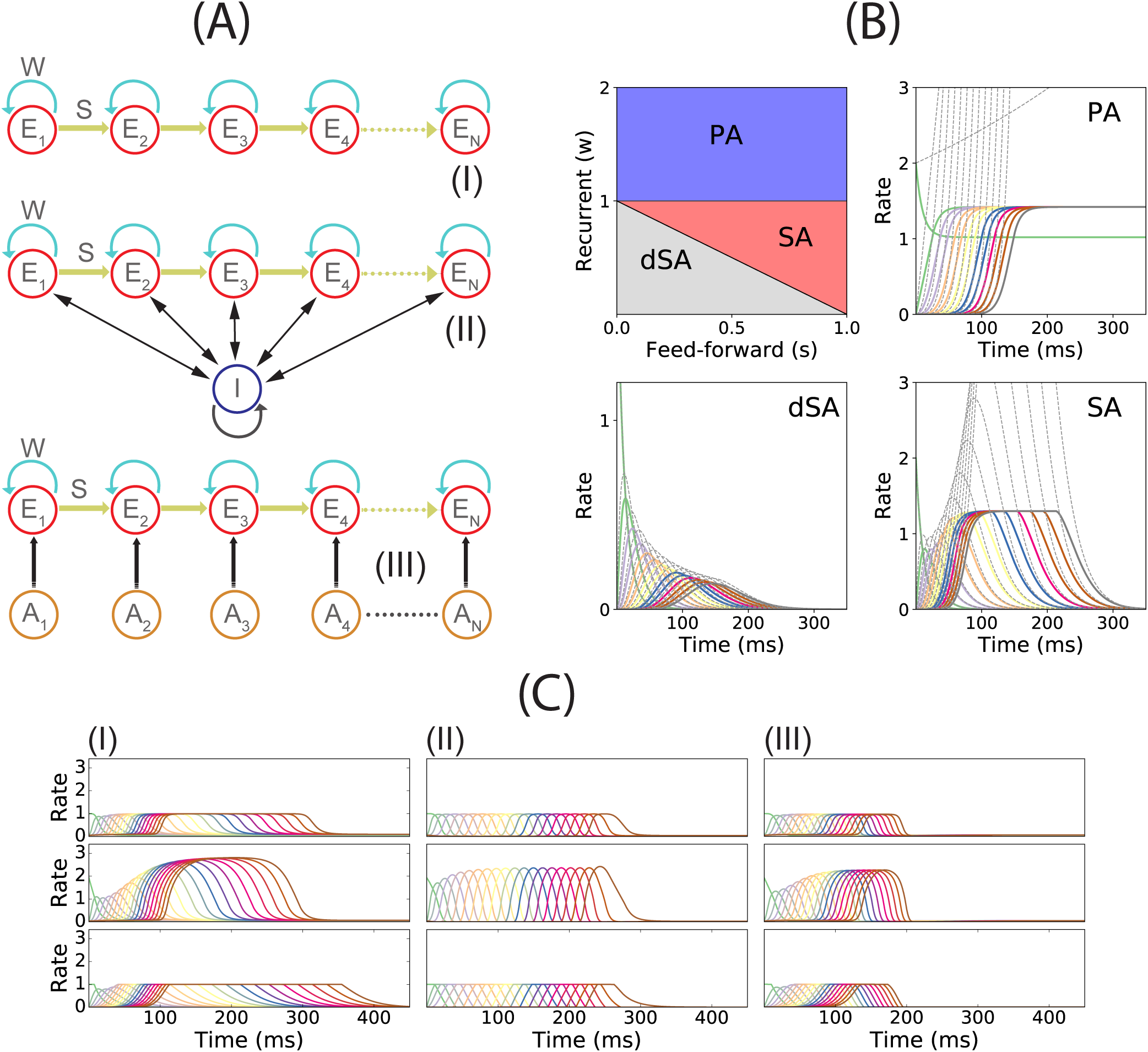
PA and SA generation in a network with fixed connectivity. **(A)** Three models of recurrent and feed-forward connected populations: (I) pure excitatory, (II) excitatory with shared inhibition and (III) excitatory with adaptation. **(B)** Phase diagram for model (I) using a piecewise linear transfer function (top-left plot) and examples of the dynamics corresponding to the three phases. Dashed lines correspond to the dynamics for the same network but using a linear transfer function. **(C)** SA generation for models (I), (II) and (III) using sigmoidal (first row), piecewise nonlinear (second row) and piecewise linear (third row) transfer functions. Parameters used in panels B,C can be found in Table S1.

#### 1.1.2 Network of excitatory neurons with shared inhibition

The network consist in *n* excitatory populations connected as in section 1.1.1, and a single inhibitory population fully connected with the excitatory populations. A schematic of the network architecture is shown in Fig 1A.II. Assuming a linear inhibitory transfer function, the dynamics of the network is given by:

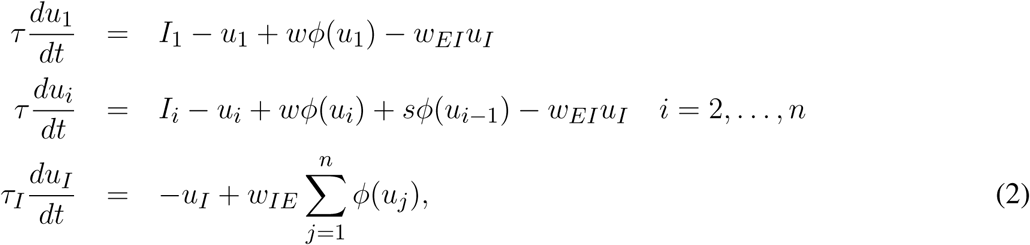

where *w*_*EI*_ is the average inhibitory synaptic strength from inhibitory to excitatory populations, *w*_*IE*_ the average inhibitory synaptic strength from excitatory to inhibitory populations and *τ*_*I*_ the characteristic time scale of the inhibitory population. When *τ*_*I*_ *« τ*, then 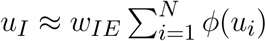 and Eq. (2) becomes

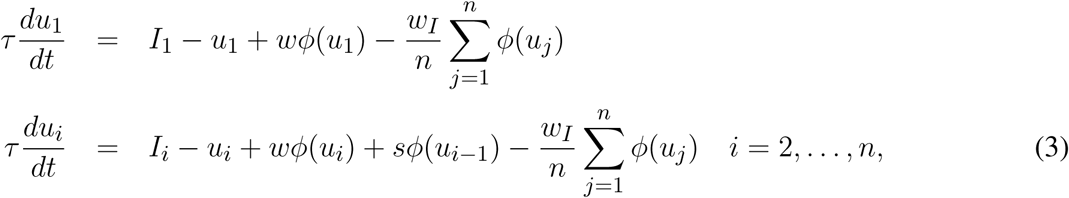

where *w*_*I*_ ≡ *nW*_*EI*_ *W*_*IE*_. See Fig. S1 in the Supplementary Material for the agreement between the full model described in Eq. (2) and its approximation in Eq. (3).

#### 1.1.3 Network of excitatory neurons with adaptation

This network consist in *n* excitatory populations connected as in sections 1.1.1 and 1.1.2 plus an adaptation mechanism for each population. A schematic of the network architecture is shown in Fig 1A.III. The dynamics of the network is given by:

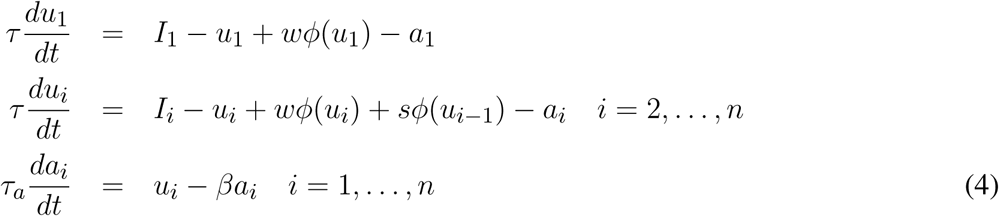

where *τ*_*a*_ is the characteristic time scale of the adaptation mechanism, and *β* measures the strength of adaptation.

### 1.2 Transfer functions

For the fixed connectivity part of this study we used three different families of transfer functions. The sigmoidal transfer function is described by

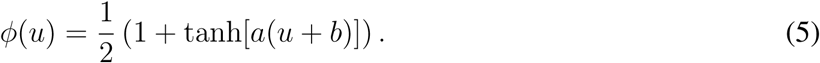

This is a saturating monotonic function of the total input, and represents a normalized firing rate. This transfer function has been widely used in many theoretical studies in neuroscience (Gerstner et al., 2014; Ermentrout and Terman, 2010), and have the advantage to be smooth. Furthermore, we have recently shown that such transfer functions provide good fits to *in vivo* data (Pereira and Brunel, 2018).

The second transfer function considered is piecewise linear:

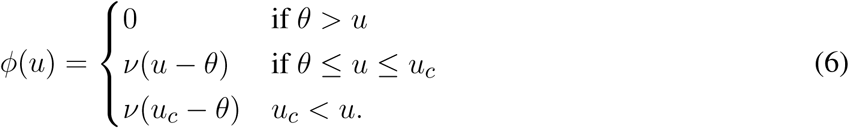

This is a piecewise linear approximation of the sigmoidal transfer function. Using this transfer function, the nonlinear dynamics of a network with a sigmoidal transfer function can be approximated and analyzed as a piecewise linear dynamical system.

The third transfer function used in this work is piecewise nonlinear (Brunel, 2003)

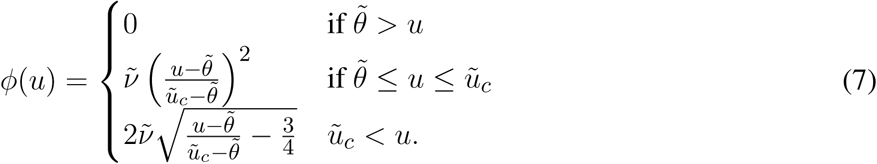

This transfer function combines several features that are present in more realistic spiking neuron models and/or real neurons: a supralinear region at low rates, described by a power law (Roxin et al., 2011), and a square root behavior at higher rates, as expected in neurons that exhibit a saddle node bifurcation to periodic firing (Ermentrout and Terman, 2010). Examples of these three transfer functions are shown in Fig 2.

**Figure 2.**
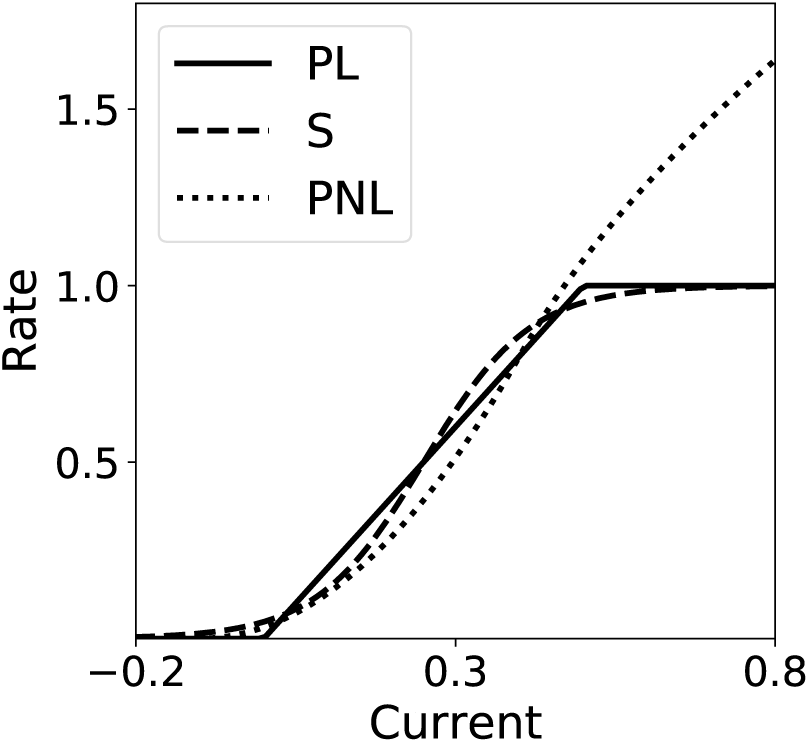
Transfer Functions. Piecewise linear (PL), sigmoidal (S) and piecewise nonlinear (PNL) transfer functions. Parameters are the same as the ones used in Fig 1.

**Figure 3.**
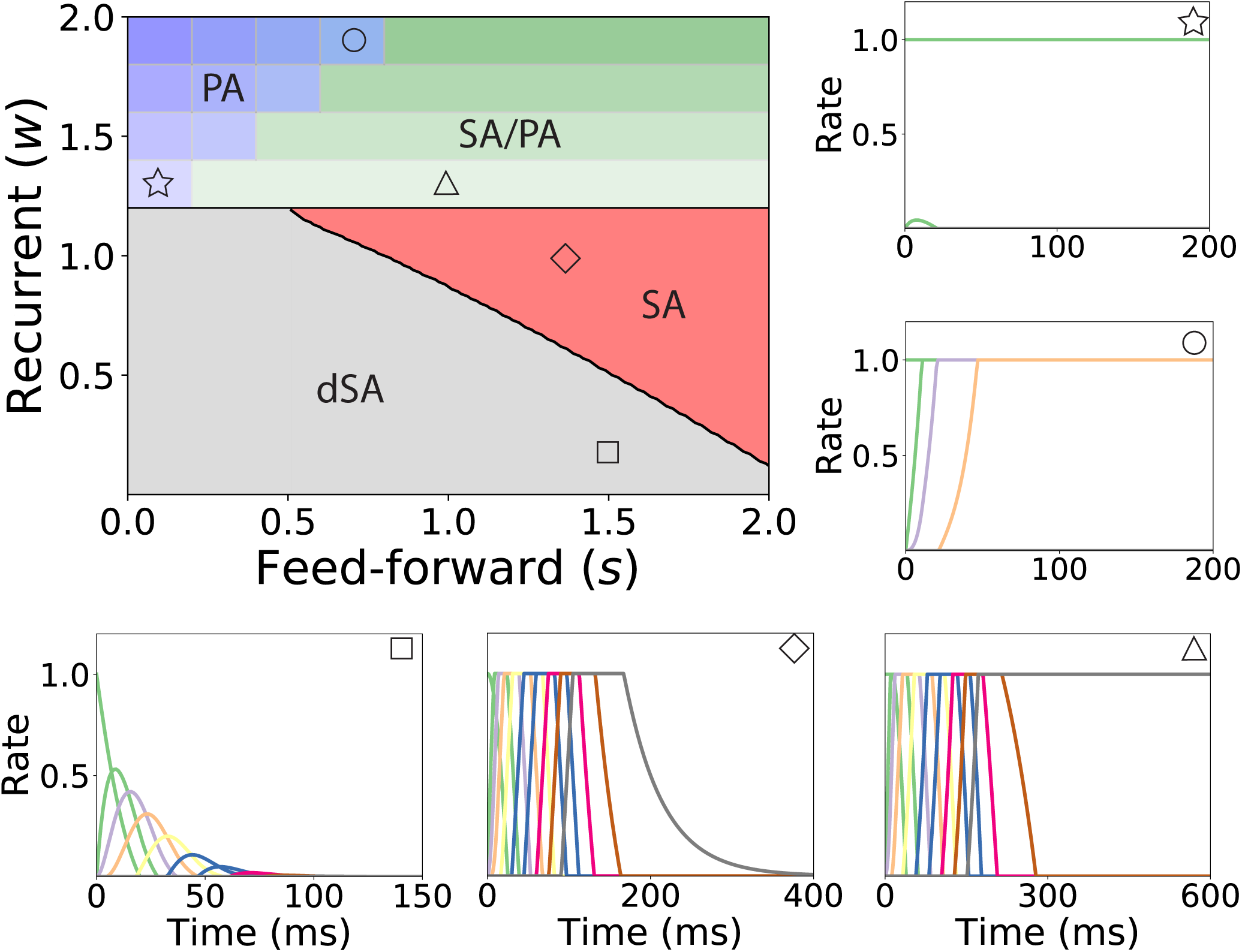
Bifurcation diagram for feed-forward-recurrent connected network of excitatory populations with shared inhibition. Top left plot: Bifurcation diagram in the *s*-*w* plane, showing qualitatively different regions: dSA (gray), SA (red), SA/PA (green) and PA (blue). The PA region is divided in sub-regions which are distingushed by the maximum and minimum number of populations active during PA (see text). The SA/PA region is also subdivided into sub-regions characterized by a different number of the maximum number of populations active in PA at the end of the sequence. Regions are separated by black lines and sub-regions are separated by gray lines. Five plots encompassing the bifurcation diagram show examples of the dynamics observed in its four qualitatively different regions. Initial condition: first population active at the maximum rate, while the rest is silent. The location in the corresponding regions of the parameter space are indicated with the symbols on the top right of the surrounding plots. Parameters can be found in Table S2.

### 1.3 Temporally asymmetric Hebbian plasticity rule

When a temporally asymmetric Hebbian plasticity rule is included (see sections 2.2-2.5 in Results), the dynamics of excitatory-to-excitatory connectivity obeys

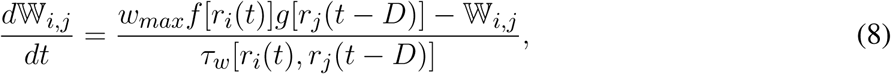

where *f* (*r*) and *g*(*r*) are sigmoidal functions given by

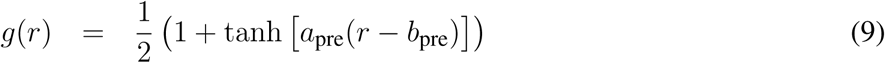

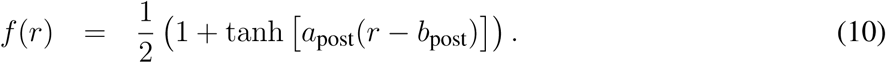

They describe the dependence of the learning rule on post and presynaptic firing rates, respectively (i.e. their dependence on *ϕ*(*u*_*i*_) and *ϕ*(*u*_*j*_)), and are bounded by zero for small or negative values of the population synaptic current, and by one for large values (see Fig 4 A and B). Here *w*_*max*_ is the maximal synaptic efficacy; *D* is a temporal delay; and *τ*_*w*_ is an activity-dependent time constant of the plasticity rule. The learning time scale is given by

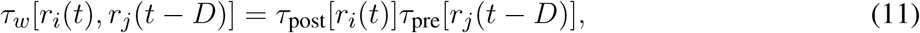

where

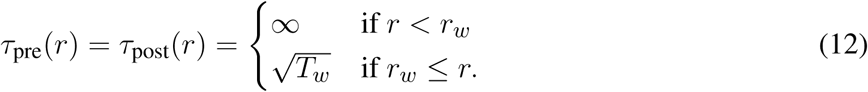

**Figure 4.**
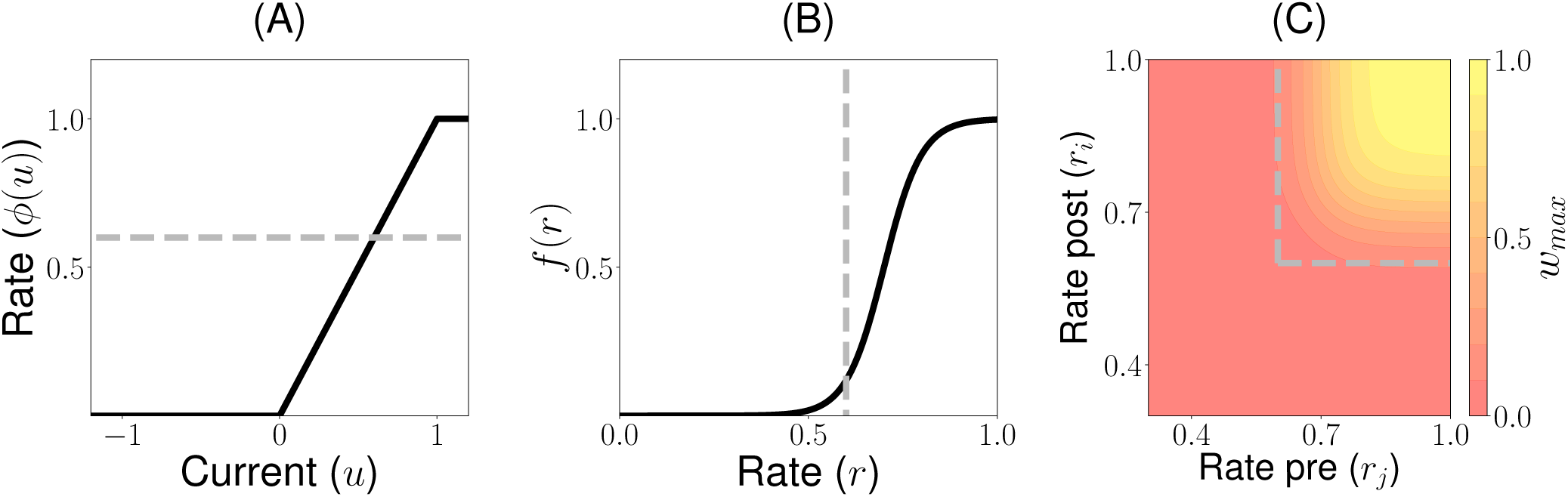
Unsupervised Hebbian learning rule: **(A)** Piecewise linear transfer function. The dashed gray horizontal line indicates the plasticity threshold *r*_*w*_. **(B)** Post synaptic dependence on the rates of the stationary connectivity function, *f* (*r*). The vertical dashed gray line indicates the plasticity threshold. **(C)** Contour plot of the stationary connectivity function, *w*_*max*_*f* (*r*_*i*_)*g*(*r*_*j*_). The dashed gray box indicates the plasticity threshold. Parameters can be found in Table S3.

Here *r*_*w*_ and *T*_*w*_ are the plasticity threshold (see dashed line in Fig 4A-C) and time scale respectively. The time scale *T*_*w*_ is chosen to be several order of magnitude slower than the population dynamics (see Table S3). When pre and/or post-synaptic currents are below a plasticity threshold *r*_*w*_, the activity-dependent time constant *τ*_*w*_ becomes infinite, and therefore no plasticity occurs. When both are above *r*_*w*_, then the activity-dependent time constant *τ*_*w*_ is equal to *T*_*w*_, and plasticity is ongoing. Thus, with this rule strong, long and/or contiguous in time enough stimuli produce lasting modifications in the synaptic weights. Otherwise, no learning occur.

### 1.4 Synaptic normalization

When a synaptic normalization mechanism is included (see section 2.3 in Results), in addition to the Hebbian plasticity rule described in section 1.3, in our network simulations, at each time step we subtracted the average synaptic change to each incoming synapse to a given neuron. This average is taken over all the incoming synapses to a particular neuron. This simulation scheme ensures that the sum of the incoming synaptic weights to each neuron remains constant, i.e.

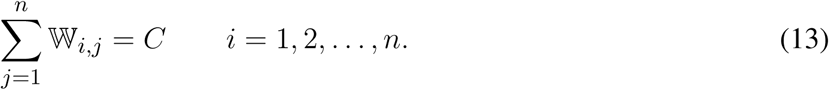

### 1.5 Multiplicative homeostatic plasticity rule

We implement a modified version of the multiplicative homeostatic rule proposed in Renart et al. (2003); Toyoizumi et al. (2014) (see sections 2.4 and 2.5 in Results). The rule is implemented in addition to the Hebbian plasticity rule described in the section 1.3. In this rule an homeostatic variable *H*_*i*_ slowly controls the firing rate of neuron *i* by scaling its synaptic weights multiplicatively. The synaptic weights will be given by

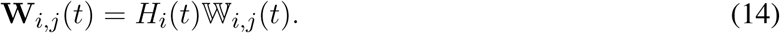

The variable 𝕎_*i,j*_(*t*) is governed by the Hebbian plasticity rule described by Eqs (8-12). The dynamics for *H*_*i*_ is given by

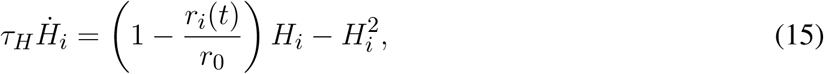

where *r*_0_ = *ϕ*(*u*_0_) is a parameter that controls the *average* firing rate of population *i* and *τ*_*H*_ is the characteristic time scale of the learning rule. Note that because of the quadratic term in the r.h.s. of Eq. (15), this rule does not in general keep the firing rates at a fixed value, and therefore this rule is not strictly speaking homeostatic. However, we keep this terminology due to the similarity with the standard homeostatic rule that does not include this quadratic term.

### 1.6 Learning dynamics under noisy stimulation

In the last section of the Results, we include noise in the population dynamics in order to asses the robustness of the learning process (see section 2.5 in Results). The equations used to describe the dynamics of the network with Hebbian and homeostatic plasticity are given by

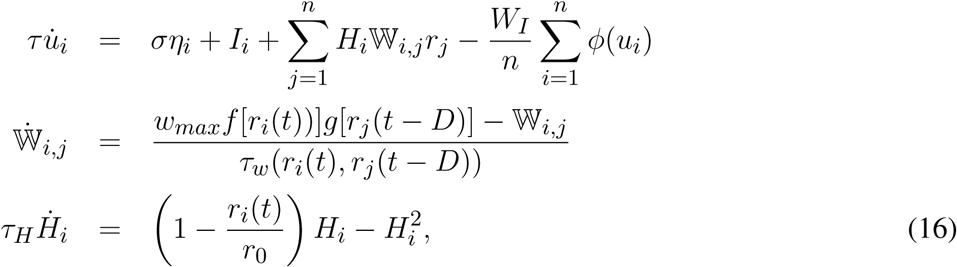

where *r*_*i*_(*t*) = *ϕ*(*u*_*i*_(*t*)) for *i* = 1, 2, *…, n* and *η*_*i*_ is a Gaussian white noise.

### 1.7 Sequential stimulation

During the learning protocol excitatory populations are stimulated sequentially once at a time for a period *T* and a time delay Δ. The stimulation can be implemented as a sequence of vectors presented to the entire the network 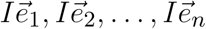, each vector corresponds to the canonical base in ℝ^*n*^ scaled by a stimulation amplitude *I*. This sequence of stimulation is repeated *k* times. To prevent a concatenation between the first and the last population stimulated, the period between each repetition *k* is much longer than *T* and Δ and any time constant of the network. Each stimulus in the sequence has the same magnitude, that is larger than the learning threshold (i.e. *r*_*w*_ < *I*). A schematic diagram of the stimulation protocol is shown in Fig 5 A.

**Figure 5.**
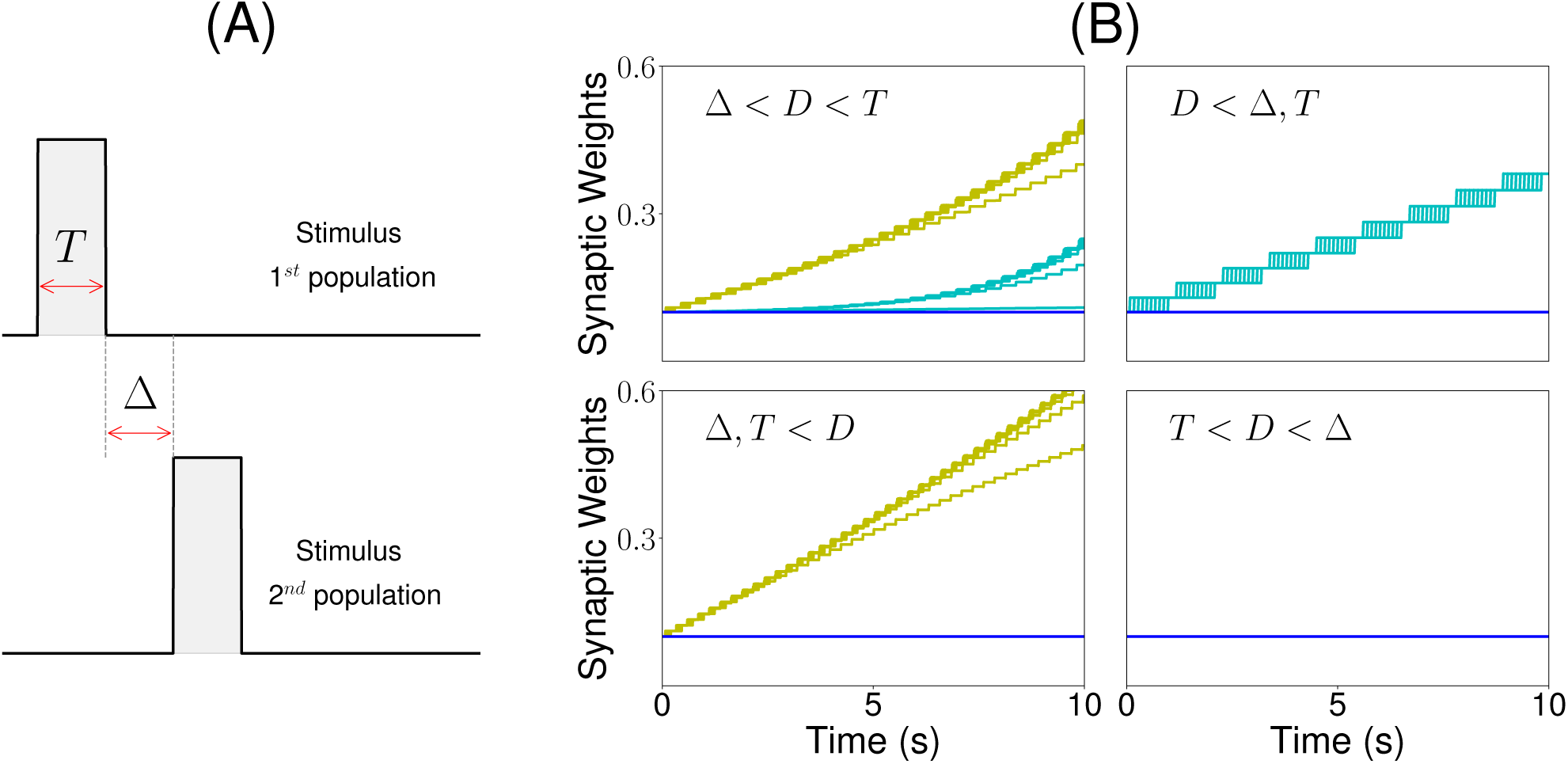
Sequential stimulation and initial synaptic weights dynamics. **(A)** Schematic diagram showing stimulation protocol for two populations. Population 1 is first stimulated for some time *T*. Then, after an inter-stimulation Δ time, population 2 is stimulated for the same duration *T*. **(B)** The weight dynamics is shown for four different stimulation regimes. Top-left: Δ < *D < T*; top-right: *D* < Δ, *T*; bottom-left: *T,* Δ < *D*; bottom-right: *T < D* < Δ. Cyan: recurrent connections; Yellow/Green: feed-forward; Blue: all other connections. Parameters can be found in Tables S3,S4.

### 1.8 Code

Simulations were performed using code written in Python. A self-contained version of the code that reproduces all the figures in this paper is available in the GitHub repository: https://github.com/ulisespereira/Unsupervised.

## 2 RESULTS

### 2.1 Persistent and sequential activity in networks with fixed connectivity

To better understand the dependence of PA and SA generation on network connectivity, we consider first a simple *n* population rate model with fixed feed-forward and recurrent connectivity (see Fig 1A). This architecture possesses the two connectivity motifs that have been classically considered the hallmarks of PA and SA — recurrent and feed-forward connections — in a space of parameters that is low dimensional enough to be suitable for full analytical treatment. In this model, the dynamics of the network is characterized by the synaptic inputs *u*_*i*_ to each population of the network (*i* = 1, *…, n*) whose dynamics obey the system of ordinary differential equations in Eq. (1). Note that we use here the *current based* formulation of the firing rate equations, that has been shown to be equivalent to the *rate based* formulation (Miller and Fumarola, 2012).

In this model, we identify the regions in the connectivity parameter space where SA, PA or decaying sequences of activity (dSA) are generated. We start with a piecewise linear transfer function with slope *v*, and compute the bifurcation diagram that gives the boundaries for qualitatively different dynamics in the parameter space (see Fig 1B and section 2 in the Supplementary Material for mathematical details). We find that robust SA can be generated provided recurrent connections are smaller than the inverse of the slope *v*, and the feed-forward connections are strong enough, *w* < 1*/? < w* + *s*. For large values of *w* (*w >* 1*/?*), the dynamics converge to a fixed point where 0*≤p≤n* populations are in a high rate state, where *p* depends on the initial conditions. When both recurrent and feed-forward connections are weak enough (i.e. *w* + *s* < 1*/?*) the activity decays to zero firing rate fixed point, after a transient in which different populations are transiently activated - a pattern which we term decaying sequence of activity or dSA.

This picture is qualitatively similar when other types of nonlinear transfer functions are used (see Methods and Fig 2 for the transfer functions used in this paper). The saturation nonlinearity of the transfer function is key to generate long lasting (non-attenuated) SA even when the number of populations is large. In a linear network, sequential activity would increase without bound for an increasing number of populations participating in the SA (see Fig 1B, dashed lines and section 2 in the Supplementary Material for mathematical details). During sequential activity, each population is active for a specific time interval. We used the analytical solution of the linearized system (see Eq. S4) to show that the duration of this active interval scales as the squared root of the position of the population along the sequence. This implies that for long lasting SA the fraction of active populations will increase with time (see Fig 1B). This feature is not consistent with experimental evidence that shows that the width of the bursts of activity along the sequence is approximately constant in time (Hahnloser et al., 2002; Harvey et al., 2012). In the model, we can prevent this phenomenon by including negative feedback mechanisms to our network architecture, either global inhibition (see Fig 1A.II) or adaptation (see Fig 1 A.III). We found that in both cases the network robustly generates PA and SA in which the fraction of active populations is approximately constant in time. These results were also qualitatively similar when different saturation nonlinearities in the transfer function were considered (see Fig 1C).

We now turn our attention to the network of excitatory neurons with global inhibition (Fig 1 A.II), since inhibition is likely to be the dominant source of negative feedback in local cortical circuits. Inhibitory interneurons are typically faster than excitatory neurons (McCormick et al., 1985). For the sake of simplicity we set the inhibitory population dynamics as instantaneous compared with the excitatory timescale. Our numerical simulations confirm that this approximation preserves all the qualitative features of the dynamics with finite inhibitory time constants, up to values of *τ*_*I*_ = 0.5*τ* (see Fig. S1 in the Supplementary Material). Using this approximation, the connectivity of the network is equivalent to a recurrent and feed-forward architecture plus a uniform matrix whose elements are *w*_*I*_ *≡ nw*_*EI*_ *w*_*IE*_. We obtained the bifurcation diagram for such a network with a piecewise linear transfer function (see section 4 in the Supplementary Material). This new bifurcation diagram shows qualitative differences with the pure excitatory network bifurcation diagram (see Fig 3). First, a qualitatively different behavior arises, where SA ends in persistent activity (region SA/PA). Second, the PA region breaks down in *n*(*n* + 1)/2 square regions of size *w*_*I*_ */n × w*_*I*_ */n*. Each region is characterized by a minimum and maximum number of populations active during PA. The lower left corner of each squared region is 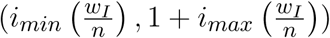 with *i*_*min*_, *i*_*max*_ = 1, 2, *…, n* (see Fig 3, different regions in graded blue), where *i*_*min*_ and *i*_*max*_ correspond to the minimum and maximum number of population active during PA within this squared region when just the first population is initialized in the active state (Fig 3 top and middle right plots). Therefore, the number of possible patterns of PA increases with the strength of the recurrent connections and decreases with strength of the feed-forward connections. On the other hand, the SA/PA is divided in *n* qualitatively different rectangular regions of size 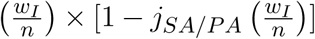 with *j*_*SA/PA*_ = 1, 2, *…, n*, where *j*_*SA/PA*_ corresponds to the number of populations that ends in PA after SA elicited by stimulating the first population in the sequence (Fig 3 bottom right plot). Then for a given strength of the recurrent connectivity *w*^*∗*^ above 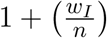, the critical feed-forward strength *s*_*c*_ that separates the PA and SA/PA regions is

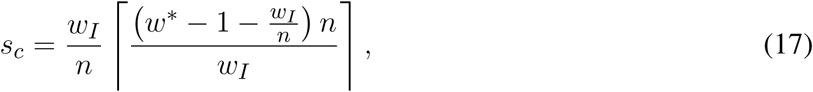

where is the ceiling function. Similarly, for a given strength of the feed-forward connection *s*^*∗*^ above 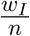, the critical recurrent strength separating SA/PA and PA is

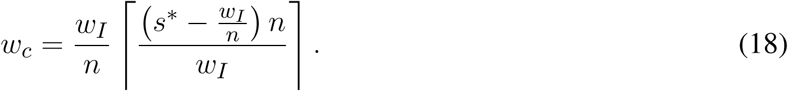

Lastly, we find that the SA region is shrunk compared with the pure excitatory network, and that the dSA region is wider.

### 2.2 Unsupervised temporally asymmetric Hebbian plasticity rule

Let us consider now a fully connected network of *n* excitatory populations with plastic synapses and global fixed inhibition. The plasticity rule for the excitatory-to-excitatory connectivity is described by Eq. (8). Using this learning rule, with fixed pre and post activity, the connectivity tends asymptotically to a separable function of the pre and post synaptic activity. The functions *f* (*r*) and *g*(*r*) are bounded by zero for small or negative values of the population synaptic current, and by one for large values (see Fig 4 A and B). This learning rule is a generalization of classic Hebbian rules like the covariance rule (Dayan and Abbott, 2001), with a non-linear dependence on both pre and post-synaptic firing rates.

The delay *D* in the learning rule leads to a temporal asymmetry (Blum and Abbott, 1996; Gerstner and Abbott, 1997; Veliz-Cuba et al., 2015). This delay describes the time it takes for calcium influx through NMDA receptors to reach its maximum (Sabatini et al., 2002; Graupner and Brunel, 2012). When this learning rule operates and the network is externally stimulated, the connectivity changes depending on the interaction of the input, the network dynamics and the learning rule. Due to the relaxational nature of Eq. (8), for long times with no external stimulation the connectivity matrix will converge to a stationary rank-1 matrix with entries of the form 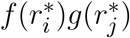, where 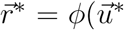 is the stationary firing rate vector, independent of all inputs presented in the past. Therefore, stimuli learned in the connectivity matrix will be erased by the background activity of the network for long times after stimulation. To prevent this inherent forgetting nature of the learning rule we introduce an activity-dependent plasticity time scale in Eqs. (11,12). Thus, when pre and/or post-synaptic currents are below a plasticity threshold *r*_*w*_, the time scale becomes infinite, and therefore no plasticity occurs. When both are above *r*_*w*_, then the time constant is given by *T*_*w*_ (see equation (12) and Fig 4). Lastly, the time scale *T*_*w*_ of these changes are chosen to be several order of magnitude slower than the population dynamics, consistent with the time it takes (∼ 1 minute or more) for plasticity to be induced in standard synaptic plasticity protocols (see e.g. Markram et al. (1997); Bi and Poo (1998); Sjöström et al. (2001), but see Bittner et al. (2017)).

Our goal is to understand the conditions for a sequential stimulation to lead the network dynamics to PA or SA, depending of the temporal characteristics of the stimulus, when this plasticity rule is introduced. Here we consider a simple stimulation protocol where each population in the network is stimulated sequentially one population at a time (see Fig 5 A). In this protocol, population 1 is first stimulated for some time *T*. Then, after an inter-stimulation time Δ, population 2 is stimulated for the same duration *T*. The other populations are then stimulated one at a time (3, 4, …, *n*) using the same protocol. The amplitude of the stimulation is fixed such that the maximum of the current elicited in each population is greater than the plasticity threshold of the learning rule. The time interval between each repetition of the sequence is much longer than *T* and Δ and any time constant of the network. When the duration of each stimulation is larger than the synaptic delay (i.e. *D < T*), recurrent connections increase, since the Hebbian term driving synaptic changes (*f* [*r*_*i*_(*t*)]*g*[*r*_*i*_(*t-D*)], where *i* is the stimulated population) becomes large after a time *D* after the onset of the presentation. When the inter-stimulation time is smaller that the synaptic delay (i.e.Δ < *D*), then the the feed-forward connections increase, since the Hebbian term driving synaptic changes (*f* [*r*_*i*+1_(*t*)]*g*[*r*_*i*_(*t − D*)]) is large in some initial interval during presentation of stimulus *i* + 1.

As a result, there are four distinct regions of interest depending on the relative values of the Δ and *T* with respect to the synaptic delay *D*. When *T* is larger than the synaptic delay, and Δ is smaller than the synaptic delay, both recurrent and feed-forward connections increase. When *T* is larger than the synaptic delay and Δ is much larger than *D*, only the recurrent connections increase. When Δ is smaller than the synaptic delay and *T* is much smaller, only the feed-forward connections increase. Lastly, when Δ is larger and *T* is smaller than *D* no changes in the connectivity are observed. The initial temporal evolution of both recurrent and feed-forward weights in representative examples of the four regions is presented in Fig 5 B. We chose not to study the region corresponding to 2*T* + Δ < *D* here, which is a region where ‘feed-forward’ connections involving non-nearest neighbor populations can also increase during learning.

We found that this learning rule is in general unstable for long sequential stimulation when both feed-forward and recurrent connections increase during the stimulation (i.e. Δ < *D < T*) to values large enough to produce persistent activity states. This is a consequence of the classic instability observed with Hebbian plasticity rules, where a positive feedback loop between the increase in synaptic connectivity and increase in firing rates leads to an explosive increase in both (Dayan and Abbott, 2001). Larger feed-forward and recurrent connections lead to an increase in number of populations active at the same time during stimulation (see Fig 6 A and D) which produce an increase of the overall connectivity by the synaptic plasticity rule (Fig 6 B and C). This leads to an increase in the overall activity producing longer periods of PA during stimulation until a fixed point where many populations have high firing rates is reached, and the connectivity increases exponentially to its maximum value (see Fig 6 B and C). By increasing the plasticity threshold, it is possible to increase the number of stimulations (and consequently the strength of the feed-forward and recurrent connections) where the network’s activity is stable. However, this does not solve the problem, since the instability on the weights eventually occurs but for a larger number of stimulations and stronger synaptic weights. In order to prevent this instability, we investigate in the next sections two different stabilization mechanisms: synaptic normalization and homeostatic plasticity. Throughout this paper, for testing whether PA, SA, SA/PA or dSA is learned, after sequential stimulation we stimulate the first population and then check whether the network recalls the corresponding type of activity (see Fig 3).

**Figure 6.**
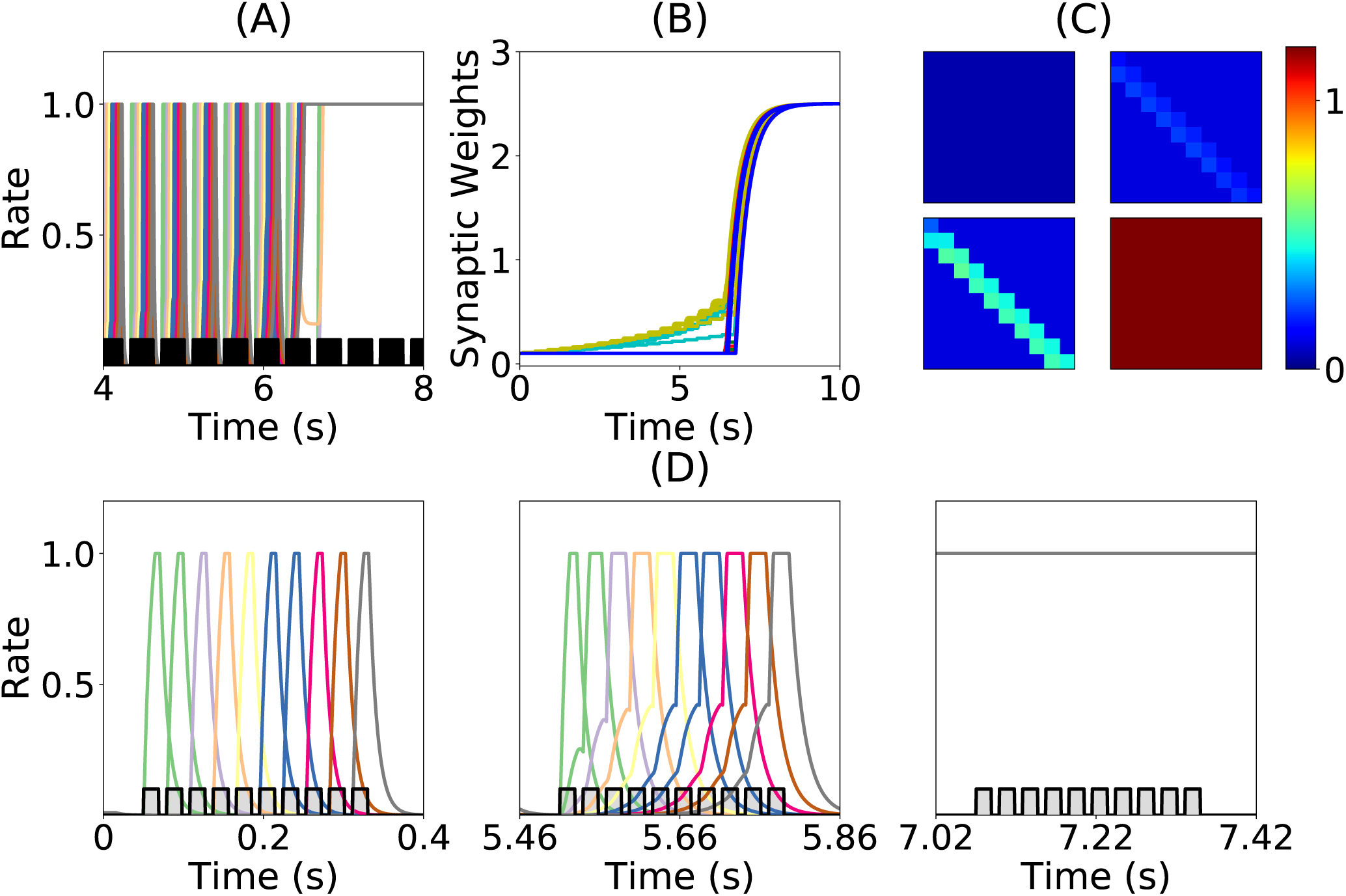
Runaway instability of the unsupervised Hebbian learning rule. **(A)** Population dynamics during 10s of sequential stimulation with *T* = 19ms and Δ = 10ms. After about 6s, all populations become active at maximal rates. **(B)** Synaptic weights dynamics during stimulation. Color code as in Fig 4D. **(C)** Connectivity matrix at different stimulation times. From left to right and from top to bottom: 0s, 3s, 6s and 9s. **(D)** Three examples of population dynamics during a single sequential stimulation at 0s, 5.46s and 7.02s respectively. Note the buildup of activity preceding each stimulus presentation because of the build-up in the feedforward connectivity at 5.46s. In A and D the black and gray traces indicate a scaled version of the stimulus. Parameters can be found in Tables S3,S4.

### 2.3 Synaptic normalization

The first mechanism we consider is synaptic normalization. This mechanism is motivated by experimental evidence of conservation of total synaptic weight in neurons (Royer and Paré, 2003; Bourne and Harris, 2011). In our model, we enforce that the sum of the incoming synaptic weights to a given population is fixed throughout the dynamics (see Eq. 13 in Methods). This constraint prevents the growth of all the synaptic weights to their maximum value during sequential stimulation due to the Hebbian plasticity, as is described in the previous section. This leads to an heterogeneous dynamics in the synaptic weights where they strongly fluctuate in time during the stimulation period, see Fig 7B. We find that the network does not reach a stable connectivity structure, and that the connectivity after the stimulation markedly depends on the specific moment when stimulation ended for a large range of stimulation parameters.

**Figure 7.**
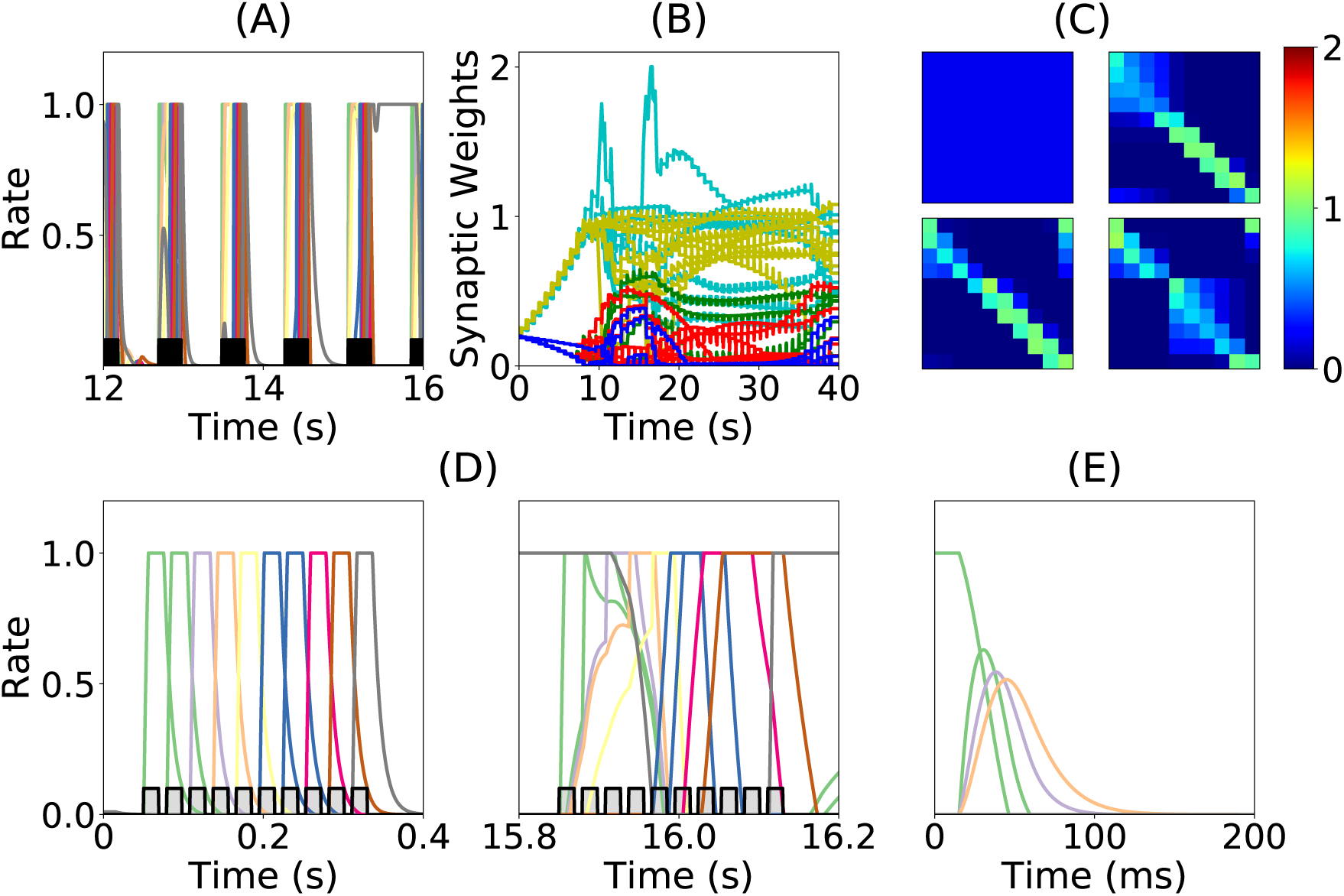
Heterogeneous synaptic dynamics for Hebbian plasticity and synaptic normalization. **(A)** Population dynamics during 10s of sequential stimulation with *T* = 19ms and Δ = 10ms. **(B)** Synaptic weights dynamics during stimulation. Cyan: recurrent connections; Light Yellow/Green: feed-forward; Red: feed-backward; Blue: feed-second-forward; Green: feed-second-backward. **(C)** Connectivity matrix at different stimulation times. From left to right and from top to bottom: 0s, 13.8s, 27.6s and 41.5s. **(D)** Two examples of population dynamics during a single sequential stimulation at 0s and 15.8s respectively. In A and D the black and gray traces indicate a scaled version of the stimulus. **(E)** Network dynamics after learning for the initial condition where the first population is active at high rate and the rest silent. Parameters can be found in Tables S3,S4.

At the initial stages of the stimulation, feed-forward and recurrent connections grow, while the rest of the synaptic connections decrease at the same rate (see Fig 7 B). When the feed-forward and recurrent connections are large enough for producing persistent activity, co-activation between a population(s) undergoing persistent activity and the population active due to the stimulation (which are not necessarily adjacent in the stimulation sequence, see Fig 7A,D) produce an increase in feed-back and upper triangular connections that are different than feed-forward and recurrent (see Fig 7B). In turn, feed-forward and recurrent connections decrease due to the synaptic normalization mechanism. This leads to complex dynamics in the synaptic weights, in which the connections sustaining co-active neuronal assemblies learned via Hebbian plasticity are depressed due to the interplay between synaptic normalization and sequential stimulation. This then leads to the formation of new assemblies due to the interplay of Hebbian plasticity and sequential stimulation.

During stimulation, the feed-forward and recurrent connectivity studied in the first section increase first, leading then in a second stage to clustered connectivities with strong bi-directional connections (see Fig 7C). Therefore, neither persistent nor sequential activity can be learned consistently after long times (see Fig 7E). Moreover, it is not clear whether neural circuits can use the observed complex synaptic dynamics to store retrievable information about the external stimuli. Thus, we find that synaptic normalization is not sufficient in this case to stabilize learning dynamics and to lead to a consistent retrieval of PA or SA. We checked that this finding is robust to changes in parameters, in particular the sum of incoming synaptic weights. In the next section we consider a second stabilization mechanism, namely Homeostatic plasticity.

### 2.4 Multiplicative homeostatic plasticity

Homeostatic plasticity is another potential stabilization mechanism that has been characterized extensively in experiments (Turrigiano et al., 1998; Turrigiano, 2017). The interplay between homeostatic plasticity and Hebbian plasticity has recently been the focus of multiple theoreotical studies (Renart et al., 2003; Toyoizumi et al., 2014; Keck et al., 2017). Here, we study the effect of multiplicative homeostatic and Hebbian plasticity for learning SA and PA. We consider a model for homeostatic plasticity in which the overall connectivity at each time **W**_*i,j*_(*t*) is given by the multiplication of two synaptic variables with different time scales as is shown in Eq. (14). In this equation, the fast plastic variable 𝕎_*i,j*_(*t*) (time scale of seconds) is governed by Hebbian plasticity, see Eq. (8). On the other hand, the slow (with a time scale of tens to hundred of seconds) homeostatic variable *H*_*i*_(*t*) scales the incoming weights to population *i*, ensuring that the network maintain low average firing rates on long time scales. Its dynamics of the homeostatic variable is given by Eq. (15). This is a modification of the standard homeostatic learning rule (Renart et al., 2003; Toyoizumi et al., 2014), that does not include the quadratic term in the r.h.s. of Eq. (15). The equation proposed in (Toyoizumi et al., 2014) stabilizes the network’s activity during stimulation, preventing the runaway of the firing rates and synaptic weights. Scaling down the overall connectivity during stimulation prevents co-activation of multiple populations, and lead to stable learning, see Fig S2D and E. However, in the network’s steady state (i.e. when times longer than the time scale of the homeostatic variable have passed without any stimulation), if the equation proposed in (Toyoizumi et al., 2014) is used, then each connection will be proportional to the factor 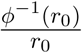 multiplied by a number of order one (see section 5.1 and 5.2 in the Supplementary Material for a general discussion and the corresponding mathematical details respectively). This implies that the steady state connectivity after learning will depend sensitively on the choice of the value of the objective background firing rate (i.e. *r*_0_) and the specific functional form of the transfer function (i.e. *ϕ*(*u*)). Due to the transfer function nonlinearity, small changes in *r*_0_ might produce large values for the factor 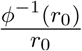 and therefore very strong connections for the steady state connectivity (see Fig S2). This is due to the fact that steady state large values in the homeostatic variable *H* scale up the connectivity learned via Hebbian plasticity in a multiplicative fashion, see Eq. (14). In practice, PA is retrieved almost always independently of the type of stimulation presented during learning, and in the absence of the quadratic term in Eq. (15) no temporal attractor other than PA can be learned. This problem can be prevented by the introduction of a quadratic term in the original homeostatic rule (see section 5 in the Supplementary Material). Note that with this quadratic term, the homeostatic plasticity rule does not exactly achieve a given target firing rate, and therefore is not strictly speaking ‘homeostatic’. However, since it is variant of the classic linear homeostatic rule, we have chosen to stick with this terminology.

We explore the role of this multiplicative homeostatic learning rule for learning both PA and SA. During sequential stimulation, the average firing rate is higher than the background objective firing rate *r*_0_, and the homeostatic variables decrease to values that are smaller than 1, see Fig 8 A and C. As a result, during sequential stimulation the dynamics of the homeostatic variable will be dominated by the linear version of the homeostatic learning rule proposed in (Toyoizumi et al., 2014), since 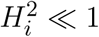 Then, the small values that the homeostatic variables take during the sequential stimulation scale down the increasing values of the recurrent and feed-forward connections due to Hebbian plasticity. This produces a weak excitatory connectivity during a repeated sequential stimulation (see Fig 8 C), preventing activation of spurious populations during stimulation (see Fig 8 B), even though the strength of recurrent and feed-forward connections learned via Hebbian plasticity are strong enough to produce PA or SA, since these connections are *masked* by the homeostatic variable. When the network returns to the steady state after sequential stimulation, the homeostatic variables return to values *H*_*i*_*∼ 𝒪* (1) (see section 5.2 in the Supplementary Material for the mathematical details), and the recurrent and feed-forward connections learned via Hebbian plasticity are *unmasked*. This mechanism stabilizes learning, allowing the network to stably learn strong recurrent and feed-forward connections, consistent with SA or PA dynamics (see Fig 8D).

**Figure 8.**
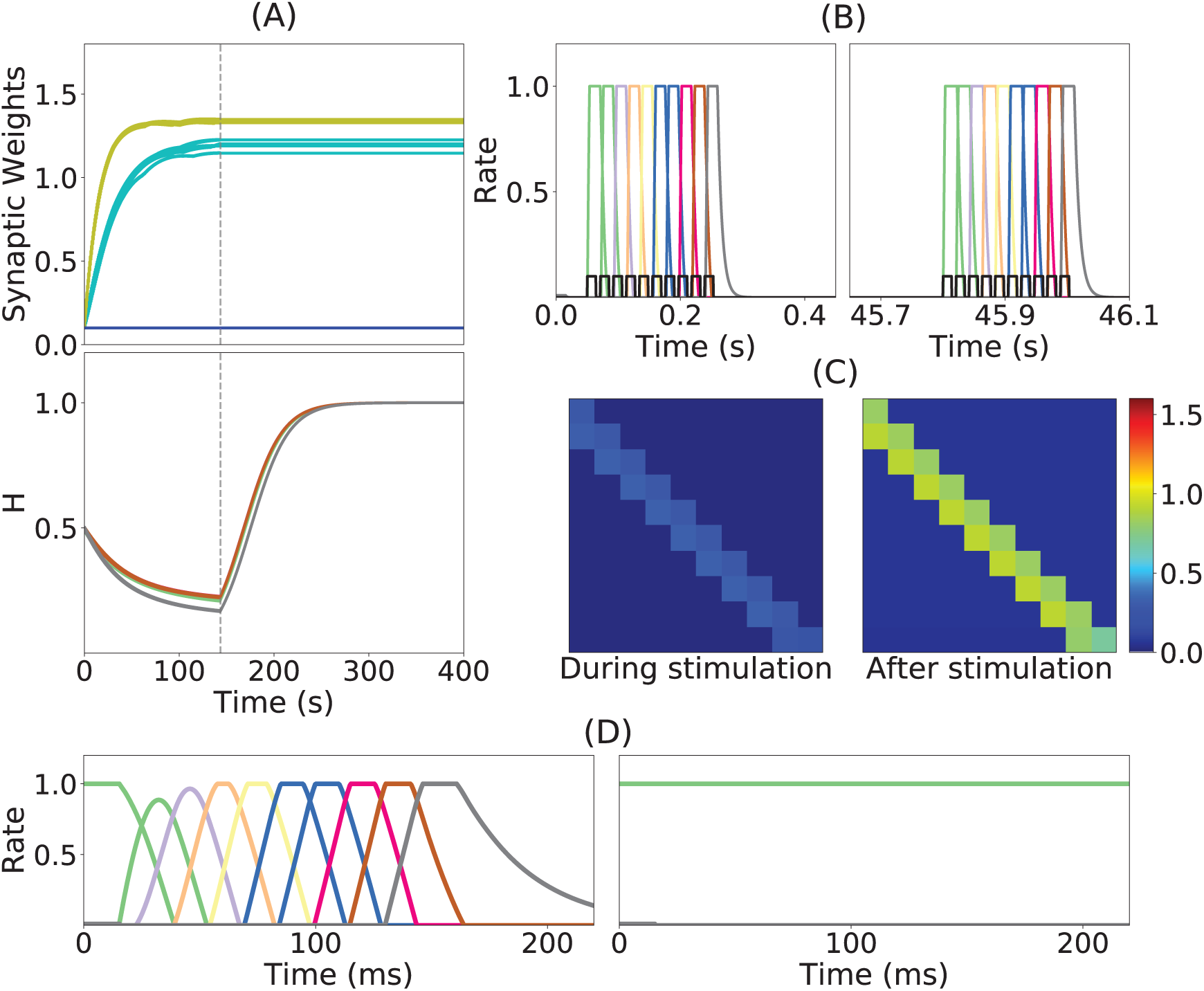
Learning dynamics in a network with Hebbian and multiplicative homeostatic plasticity: **(A)** Top: synaptic weights dynamics during and after stimulation. Cyan: recurrent; Yellow: feed-forward; Blue: all other connections. Bottom Homeostatic variables in excitatory populations. neuron *i*. Gray vertical dashed line indicate the end of the sequential stimulation. **(B)** Neuron dynamics during stimulation for two different periods of time. **(C)** Snapshots of the connectivity matrix **W**_*i,j*_(*t*) at the end of the sequential stimulation (left) and 60s after the end of the sequential stimulation (right). **(D)** Network dynamics after learning following an initial condition where the first population is active at high rate while all others are silent for two different stimulation parameters, for two stimulation parameters, one that generates SA (left), the other PA (right). Parameters can be found in Tables S3,S4.

The weakening of recurrent connections during sequential stimulation allows us to derive an approximate analytical description of the temporal evolution of the synaptic connectivity with learning. Since the net current due to connections between populations is very small, each population dynamics is well approximated by an exponential rise (decay) toward the stimulation current (background current) provided inhibition is weak enough (see Fig 9). By using this approximation we build a mapping that yields the value of the recurrent and feed-forward synaptic strengths as a function of stimulation number *k*, stimulation period, *T*, and delay, Δ (see Eqs. (S32,S33) in 6 of Supplementary Material). This mapping provides a fairly accurate match of both the dynamics of the synaptic weights and the final steady state connectivity matrix in the case of weak inhibition (see Fig 10A, corresponding to *w*_*I*_ = 1) and a less accurate match for stronger inhibition (see Fig 10B, *w*_*I*_ = 2). This is expected since our theoretical analysis neglects the effect of inhibition during learning (see section 6 of Supplementary Material). The mapping derived for evolution of the synaptic weights during sequential stimulation corresponds to a dynamical system in the (*s, w*) phase space that depends on the stimulus parameters (Δ, *T*) and the initial connectivity. The final connectivity corresponds to the fixed point of these dynamics (see Eqs. (S34,S35) in section 6 of Supplementary Material).

**Figure 9.**
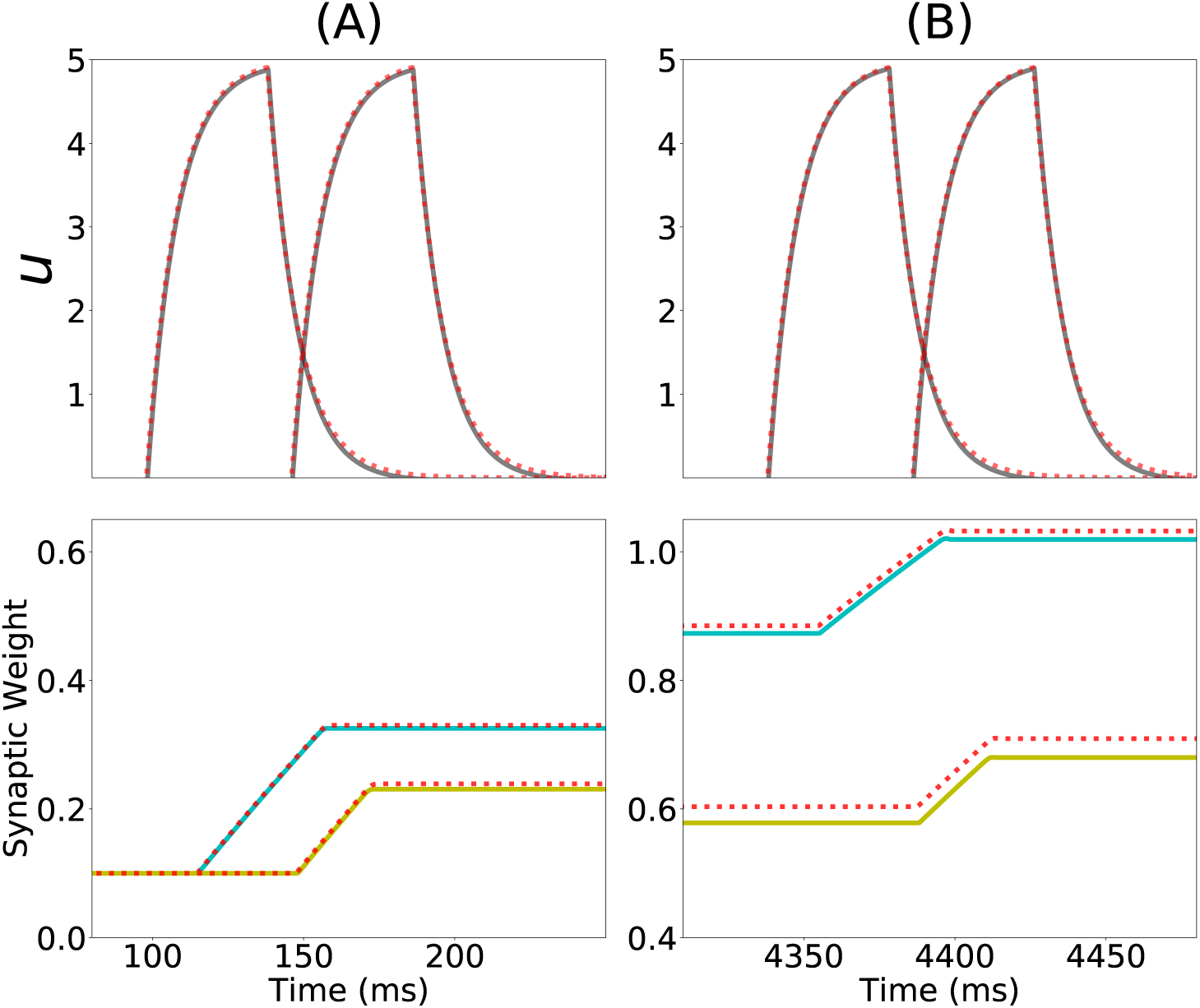
Analytical approximation of the dynamics of the network with Hebbian and multiplicative homeostatic plasticity: (**First row**) Current dynamics for the second and third populations in a network of 20 populations during one presentation of the sequence. The dashed red line shows the analytical approximation for the dynamics during stimulation (Eq. S24 in section 6 of Supplementary Material). (**Second row**) Dynamics of the recurrent synaptic strength within the second population (cyan), and the ‘feed-forward’ synaptic strength from the second to the third population (yellow) during the same presentation of the sequence. The dashed red line shows the analytical approximation for the synaptic weight dynamics (Eq. (S26,S30) in section 6 of Supplementary Materials). (**A**) and (**B**) correspond to the first and the fifth presentation of the stimulation sequence respectively. Parameters can be found in Tables S3,S4.

**Figure 10.**
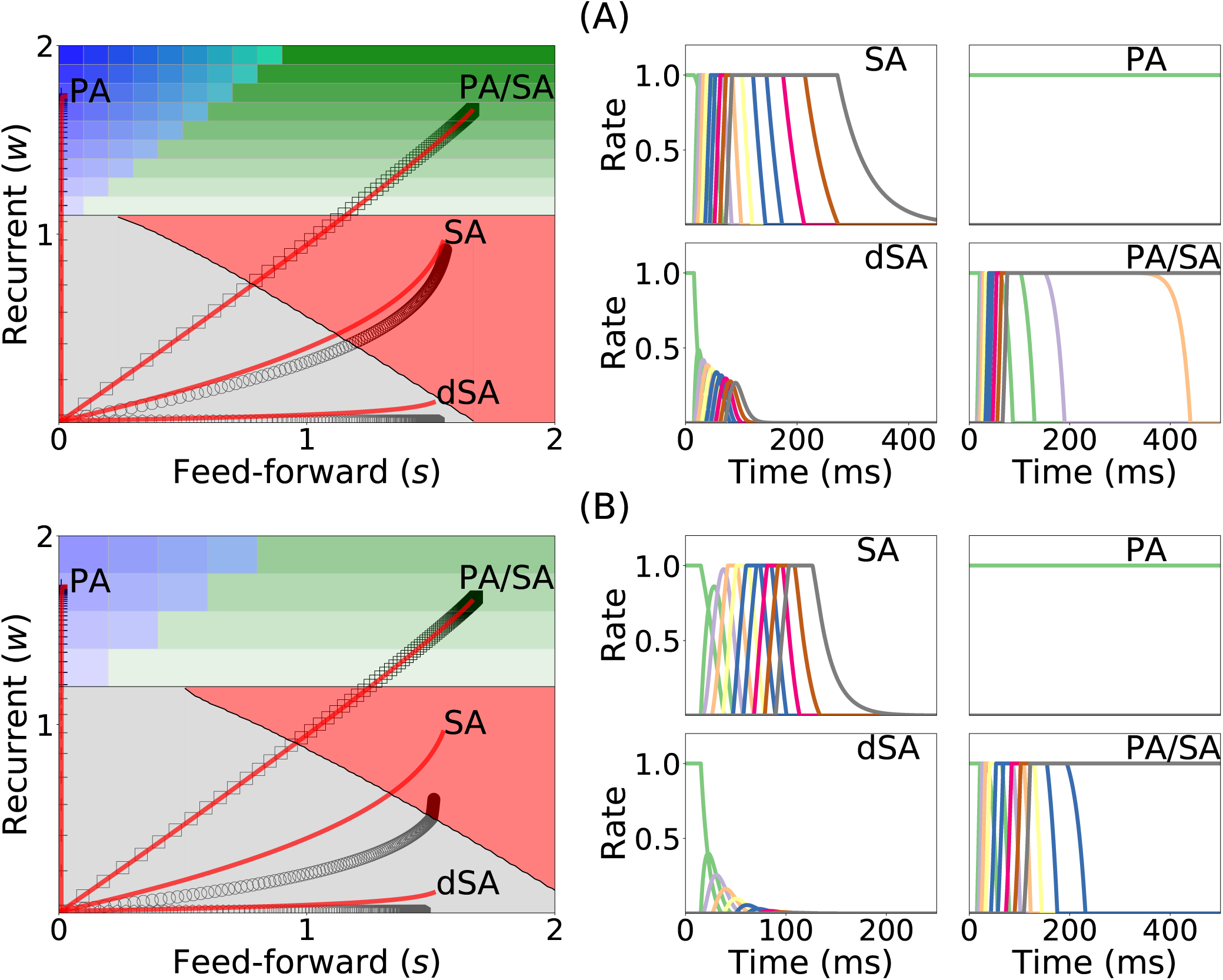
Changes in recurrent and feed-forward synaptic strengths with learning, for different sequences with different temporal parameters. (**Left**) Dynamics of recurrent and feed-forward connections in the (*s, w*) parameter space during sequential stimulation for four different values of Δ and *T*. Black circles (SA), plus signs (PA), hexagons (dSA), and squares (PA/SA) show the simulated dynamics for (*T,* Δ) ={(7, 14), (50, 40), (5, 13), (20, 8.5)} (in ms) respectively. Red traces indicate the approximated dynamics derived in section 6 of Supplementary Material. (**Right**) Rates dynamics after many presentations of the sequence. The first population was initialized at high rates, the others at low rates. (**A**) and (**B**) correspond to *w*_*I*_ = 1 and *w*_*I*_ = 2 respectively. Parameters can be found in Tables S3,S4.

Fig 10 shows that depending on the temporal characteristics of the input sequence, the network can reach any of the four qualitatively different regions of the phase diagrams in a completely unsupervised fashion. For values of Δ that are smaller than the synaptic delay *D* and *T* on the order or larger than *D*, the network generates SA. For values of *T* approximately larger than *D* and for Δ small enough, the dynamics lead to SA/PA. Lastly PA is obtained for large enough Δ and *T*. These observations match with the intuition that stimulations long enough but far delayed in time leads to learning of PA and that stimulations contiguous in time but short enough leads to SA. Stimulations between these two conditions (long and contiguous) leads to a combination of both dynamics, i.e. SA/PA, as shown in Fig 10.

### 2.5 Learning and retrieval is robust to noise

Under *in vivo* conditions neural systems operate with large amount of variability in their inputs. In order to assess the effect of highly variable synaptic input current during learning and retrieval, we add a mean zero uncorrelated white noise to the dynamics when both Hebbian learning and homeoestatic plasticity are included in the network, as described in Eq. (16). We found that both the synaptic weights dynamics during learning and the retrieved spatiotemporal dynamics after learning are robust to noise (see Fig 11), even when the amplitude of the noise is large (i.e. inputs with values equal to the standard deviation of the noise lead to a population to fire at 30% of the maximum firing rate). During sequential stimulation, the learning dynamics is marginally altered for both weak and strong inhibition (compare Fig 11 with Fig 10).

**Figure 11.**
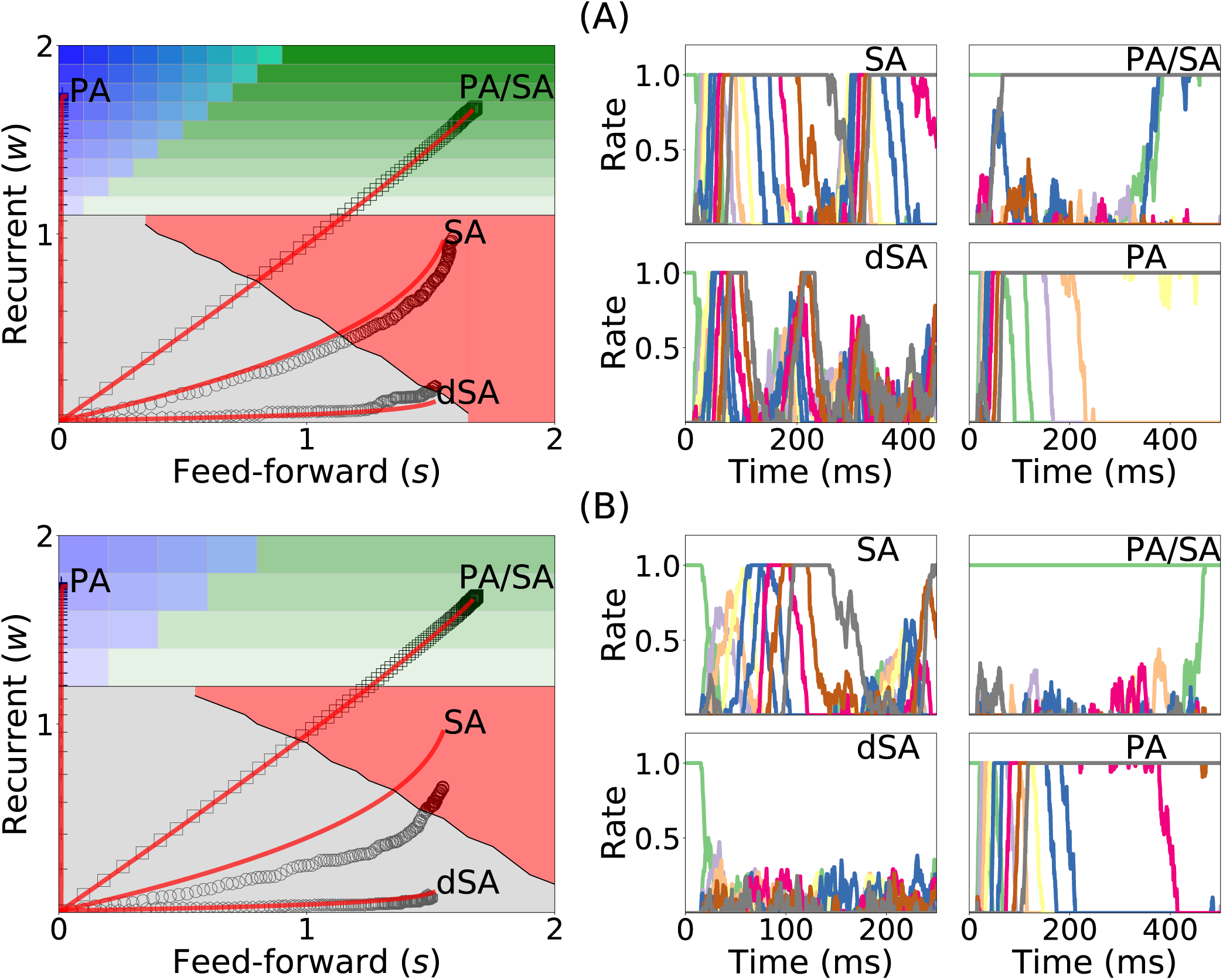
Learning dynamics under noisy stimulation. Same as in Fig 10, but in the presence of a white noise input current, with mean 0 and standard deviation of 0.3 (i.e. *σ* = 0.3 in Eq. (16)).

Importantly, the synaptic weights reach very similar stationary values compared with the case without noise. After learning, even though the rates stochastically fluctuate in time, the retrieved spatiotemporal attractors (i.e. PA, SA, dSA or PA/SA) are qualitatively similar as in the case without noise (compare Fig 11 with Fig 10). One qualitative difference in the case with external noise, is that in both SA and PA/SA dynamical regimes random inputs lead to a repetition of the full or partial learned sequence. Altogether, this simulations show that the network can robustly learn and retrieve qualitatively the same spatiotemporal attractors in the presence of external noise.

## 3 DISCUSSION

We have shown that under sequential stimulation a network with biologically plausible plasticity rules can learn both PA or SA depending on the stimulus parameters. Two plasticity mechanisms are needed: 1) Hebbian plasticity with temporal asymmetry; 2) a stabilization mechanism which prevents the runaway of synaptic weights while learning. When unsupervised Hebbian plasticity is present alone the network fails to stably learn PA or SA, while including multiplicative homeostatic plasticity stabilizes learning. For stable learning, we show that the learning process is described by a low dimensional autonomous dynamical system in the space of connectivities, leading to a simplified description of unsupervised learning of PA and SA by the network from external stimuli. Depending on the stimulus parameters, the network is flexible enough to learn selectively both types of activity by repeated exposure to a sequence of stimuli, without need for supervision. This suggests that cortical circuits endowed with a single learning rule can learn qualitatively different neural dynamics (i.e. persistent vs sequential activity) depending on the stimuli statistics.

Using the full characterization of the bifurcation diagram in the space of fixed feed-forward and recurrent connections developed here, we mapped the evolution of the connectivity during stimulation in the bifurcation diagram. We analytically and numerically showed that the synaptic weights evolve in the feed-forward–recurrent synaptic connections space until they reach their steady state (when the number of sequential stimulations is large). The specific point of the steady state in the bifurcation diagram depends solely on the stimulation parameters — stimulation period *T* and time delay Δ– and the connectivity initial conditions. We found that stimulations with long durations and large delays generically leads to the formation of PA, whereas stimulations with long enough durations and short delays leads to the formation of SA. Thus, persistent stimulation leads to persistent activity while sequential stimulation leads to sequential activity.

### 3.1 Learning of sequences in networks

A growing number of network models have been shown to be able to learn sequential activity. Models with supervised learning can reproduce perfectly target sequences through minimization of a suitable error function (Sussillo and Abbott, 2009; Memmesheimer et al., 2014; Laje and Buonomano, 2013; Rajan et al., 2016), but the corresponding learning rules are not biophysically realistic.

Other investigators have studied how unsupervised learning rules leads to sequence generation. Early models of networks of binary neurons showed how various prescriptions for incorporating input sequences in the connectivity matrix can lead to sequence generation (see Kuhn and van Hemmen (1991)) - or, sometimes, both sequence generation or fixed point attractors depending on the inputs (Herz et al., 1988). The drawback of these models is that they separated a learning phase in which recurrent dynamics was shut down in order to form the synaptic connectivity matrix, and a retrieval phase in which the connectivity matrix does not change anymore.

Our model removes this artificial separation, since both plasticity rule and recurrent dynamics operate continuously, both during learning and recall. However, we found that there needs to be a mechanism to attenuate recurrent dynamics during learning for it to be stable. The mechanism we propose rely on a modified version of a standard homeostatic rule. Other mechanisms have been proposed, such as neuromodulators that would change the balance between recurrent and external inputs during presentation of behaviorally relevant stimuli (Hasselmo, 2006).

The cost of not having supervision is that the network can only learn the temporal order of the presented stimuli, but not their precise timing. Veliz-Cuba et al (Veliz-Cuba et al., 2015) have recently provided a model which bear strong similarities with our model (rate model with unsupervised temporally asymmetric Hebbian plasticity rule), but includes in addition a short-term facilitation mechanism that allows the network to learn both order and precise timing of a sequence presented in input. However, their mechanisms requires precise fine tuning of parameters.

Models with temporally asymmetric Hebbian plasticity have also been investigated in the context of the hippocampus (Abbott and Blum, 1996; Gerstner and Abbott, 1997; Mehta et al., 1997; Jahnke et al., 2015; Chenkov et al., 2017; Theodoni et al., 2017). In such models, feed-forward connectivity is learned through multiple visits of neighboring place fields, and sequential activity (‘replays’) can be triggered using appropriate inputs mimicking sharp-wave ripples. Other models use unsupervised Hebbian plasticity but qualitatively distinct mechanisms to generate sequential activity. In particular, several studies showed that sequences can be generated spontaneously from unstructured input noise (Fiete et al., 2010; Okubo et al., 2015). Murray and Escola (Murray et al., 2017) showed that sequences can be generated in networks of inhibitory neurons with anti-Hebbian plasticity, and proposed that this mechanism is at work in the striatum.

### 3.2 Stabilization mechanisms

Consistent with many previous studies (Dayan and Abbott, 2001), we have shown that a network with unsupervised Hebbian plasticity under sequential stimulation leads to a runaway of the synaptic weights. This instability is due to a positive feed-back loop generated by the progressive increase of network activity leading to a progressive increase in average synaptic strength when PA or SA are being learned. One possible solution for this problem was first proposed in the context of attractor neural network models (Amit et al., 1985; Amit and Fusi, 1994; Tsodyks and Feigel’Man, 1988). In these models, patterns are learned upon presentation during a *learning phase* where synapses are plastic but there is no ongoing network dynamics. After the *learning phase*, the learning of attractors is tested in a *retrieval phase*, where the network dynamics is ongoing but synaptic plasticity is not present. Therefore, by compartmentalizing in time dynamics and learning, the network dynamics does not lead to changes in the synaptic weights during retrieval, and conversely, changes in synaptic weights do not lead to changes in the dynamics during learning. This separation prevents the observed runaway of the synaptic weights due to unsupervised Hebbian plasticity.

However, it is unclear whether such compartmentalization exists in cortical networks. In this work, we explored the alternative scenario, in which both plasticity and dynamics happen concurrently during learning and retrieval (see also Mongillo et al. (2005); Litwin-Kumar and Doiron (2014); Zenke et al. (2015) for a similar approach in networks of spiking neurons). We found that adding multiplicative homeostatic plasticity to unsupervised Hebbian plasticity leads to stable learning of PA and SA. During sequential stimulation, the increase in co-activation between multiple populations due to recurrent and feed-forward connections learned via unsupervised Hebbian plasticity is prevented by suppressing its effect in the network dynamics. Homeostatic plasticity scales down the overall connectivity producing a weakly connected network. PA and SA is prevented to occur during stimulation, which weakens the positive feed-back loop generated by the increase in co-activations of neuronal populations. After learning, the dynamic variables of the Homeostatic plasticity rule reach a steady state with values similar of what they where before stimulation (see Fig 8 A) and the connectivity learned via unsupervised Hebbian plasticity can lead to retrieval of PA and SA upon stimulation (see Fig 8 C). The homeostatic variable reaches its steady state at a value close to one, and the connectivity recovers, *unmasking* the feed-forward and recurrent learned architecture. We have also tried other stabilization mechanisms such as inhibitory to excitatory plasticity (Vogels et al., 2011) instead of homeostatic plasticity. In this case we found that stable learning of PA and SA is possible, but for distinct sets of network and stimulation parameters (data not shown).

As explained in Zenke and Gerstner (2017); Zenke et al. (2017), in order to prevent the runaway of the synaptic weights produced by Hebbian plasticity, the time-scale of any compensatory mechanism should be of the same order or faster than the Hebbian time-scale. For multiplicative homeostatic plasticity, the time-scale of the homeostatic variable *H*_*i*_ is dependent on the firing rate of neuron *i* and the target firing rate (i.e. *ϕ*(*u*_*i*_)*/ ϕ*(*u*_0_)). When the network firing rate is close to the target firing rate the homeostatic learning rule is slow, and the homeostatic mechanism seldom play a role in the dynamics. On the other hand, for high firing rates the homeostatic plasticity time-scale becomes faster, preventing the runaway of the synaptic weights. There is currently an ongoing debate about whether the time-scales of compensatory processes used in theoretical studies, as the ones used here, are consistent with experimental evidence (see e.g. Zenke and Gerstner (2017); Zenke et al. (2017)).

## AUTHOR CONTRIBUTIONS

U.P. and N.B. designed the research. U.P. and N.B. performed the research. U.P. and N.B. wrote the manuscript.

## ACKNOWLEDGEMENTS

U.P. thanks to the Champaign Public Library for providing the physical space where part of this work has been completed.

## SUPPLEMENTAL DATA

The Supplementary Material for this article can be found online at:

